# B cell subsets have different capacities for phagocytosis and subsequent presentation of antigen to cognate T cells

**DOI:** 10.1101/2025.01.28.635276

**Authors:** Daniel M Morelli, Morgan Langille, Richard Zhang, Heather C Craig, Parisa Shooshtari, Bryan Heit, Steven M Kerfoot

## Abstract

B cells have been shown to be phagocytic under some circumstances. However, the phagocytic capacity of different B cell subsets and how this is linked to Antigen (Ag) presentation or other functions has not been characterized. To address this, we developed 2 µm phagocytic Ag conjugated bead targets that target phagocytic pathways including the BCR, scavenger, Fc, and complement receptors to study potential pathways by which B cells phagocytose both cognate and non-cognate Ags. We found that while follicular B2 (Fo B), marginal zone, and B1 B cells are highly phagocytic of BCR-engaging targets through their BCR, only peritoneal cavity B1 cells could uptake non-cognate Ag-coated beads or bacteria. Despite this, B1 cells were not effective at presenting Ag to activate cognate T cells or at killing phagocytosed bacteria. Finally, analysis of scRNA-seq data revealed that these differences in phagocytic capacity could not be explained by differential expression of relevant phagocytic receptors, implying that there is likely some form of regulation in place preventing non-cognate Ag uptake by Fo B cells. Our work will help contribute to a better understanding of non-classical Ag uptake mechanisms employed by B cells and their relevance to inflammation.

## Introduction

B cells are one of the three classic professional antigen presenting cells (APCs) as defined by their baseline expression of both MHC class I and II and expression of costimulatory molecules CD80 and CD86 with activation, resulting in their ability to activate naïve Antigen (Ag)-specific T cells *ex vivo*^1,2,3^. *In vivo*, however, the activation of naïve T cells in lymphatic tissue is largely the role of dendritic cells (DCs)^4,5^, while macrophages may play a greater role in Ag presentation to T effector cells recruited to peripheral tissues^6,7^. While B cells may present Ag to T cells in several scenarios, it is best understood in the context of T cell-dependent antibody responses where follicular B (Fo B) cells present Ag internalized through their BCR to already activated T cells specific for the same Ag^1,2,3^. These interactions are the foundation of the Germinal Center response and essential for the production of high affinity antibodies^8,9^. In addition to Fo B cells, splenic marginal zone (MZ) and peritoneal cavity (PerC) B1 B cells participate in immune responses against blood pathogens^10^ and mucosal pathogens^11^ respectively, and tend to be activated independent of T cell help^3^. It is not well understood how the different functions of these subsets influence their ability to acquire and present Ag.

Professional APCs, appropriate to their biological role, possess multiple pathways through which to acquire Ag for subsequent processing and presentation to T cells. Indeed, DCs acquire Ag through non-specific mechanisms such as pinocytosis and micropinocytosis^12,13^. Both macrophages and DCs internalize potential Ags through receptor-mediated processes, including phagocytosis of larger (>0.5 µm) targets opsonized by complement proteins, antibodies, or can be recognized *via* relatively non-specific scavenger receptors^14,15,16^. B cells, on the other hand, are typically not thought to possess the same breadth of Ag uptake pathways, which are thought to be largely limited to the BCR^17^. Further, B cells are classically described as non-phagocytic^18,19^, although the original experimental source(s) of this contention is elusive and it is not clear if it is the result of generalizations across cell subsets or specific conditions.

More recent reports have challenged the idea that B cells cannot phagocytose larger Ag targets. Li *et.al.* demonstrated that B cells from fish were capable of phagocytosing beads^20^. Several studies followed that showed B cell phagocytosis remained evolutionarily conserved in several classes of vertebrates including fish, amphibians, and reptiles^6,22,23^. The authors of these studies suggested that this “ancient” function was lost in mammalian B cells^21,22,23^. However, Gao *et.al.* and Parra *et.al*. found that mouse peritoneal cavity B1 cells were capable of phagocytosing cognate Ag-coated beads and bacteria^24,25^, concluding that these “innate-like” peritoneal B1 cells are evolutionary reminiscent of the phagocytic B cells in other vertebrates^25^. However, Martínez-Riaño *et.al.* more recently challenged this paradigm as they demonstrated that mouse Fo B cells could phagocytose Ag-coated beads bound by BCR for subsequent presentation to T cells^26^.

Therefore, it is clear that B cells possess broader capacity for Ag uptake than previously appreciated. Although recent literature provides evidence of B cell phagocytosis, there are numerous discrepancies between these studies with respect to the B cell subsets and phagocytic pathways investigated, and therefore the Ag uptake and phagocytic capacity of different B cell subsets and how these capacities align with their unique functions within the immune response are not clear. Here, we performed a detailed analysis of multiple B cell subsets and found that, while Fo B, MZ, and B1 cells are highly phagocytic of BCR-engaging targets, only peritoneal cavity B1 cells could uptake non-cognate Ag-coated beads or bacteria. Despite this, B1 cells were not effective at presenting Ag to activate cognate T cells or at killing phagocytosed bacteria. In contrast, Fo B cells bound but did not internalize opsonized or other targets through non-BCR phagocytosis pathways, but very efficiently presented phagocytosed Ag to T cells.

## Materials and Methods

### Mice

C57Bl/6 and OTII TCR-transgenic [4194; Tg(TcraTcrb)425Cbn/J] mice were purchased from Jackson Laboratories. B1-8 mice^27^ with a homozygous deletion of the Jκ locus^28^ were a generous gift from Dr. A. Haberman. IgH^MOG^ MOG-sp. BCR knockin mice^29^ were received as a gift from Dr. H Wekerle. Mice expressing enhanced GFP within all nucleated cells *via* the ubiquitin promoter [4353; Tg(UBCGFP)30Scha/J], were obtained from the Jackson Laboratory. Mice were housed in a specific pathogen-free barrier at West Valley Barrier. Healthy mice between the ages of 6 and 12 weeks were used in experiments. Mice were age and sex matched across groups of an experiment. Both male and female mice were used with no apparent differences between the two sexes (*data not shown*). All animal experiments were conducted in compliance with the protocol (2019-123, 2023-145) approved by the Western University Animal Use Subcommittee in Canada.

### Antibodies and Reagents

Reagents for bead conjugation: Goat anti-mouse IgM F(ab’)_2_ was purchased from Jackson Laboratories (115-005-020), goat anti-human IgM F(ab’)_2_ from Jackson Laboratories (109-006-129), mouse C3b was purchased from CompTech (M114), mouse C3d was purchased from R&D Systems (2655-C3-050), and biotinylated PE was purchased from Cayman Chemicals (25724). Phosphocholine (860320P), phosphoserine (840034C), and rhodamine PE (810158P) was purchased from Avanti. Rat IgG (l4131) and ovalbumin (9006-59-1) was purchased from Sigma-Aldrich. Ovalbumin was biotinylated in-house using EZ-Link NHS-LC-Biotin (21336; ThermoFisher) according to the manufacturer’s protocol.

Reagents for microscopy and Flow Cytometry: The following Abs were purchased from BD Biosciences: anti-mouse CD45R A647 (RA3-6B2), anti-human CD43 BV421 (1G10), anti-CD27 human BV711 (L128). The following Abs were purchased from BioLegend: anti-CD4 mouse PerCP (RM4-5), anti-MHCII A700 (M5/114.15.2), anti-mouse CD44 FITC (IM7), anti-mouse F4/80 A700 (BM8), anti-mouse CD21 FITC (CR2/CR1), anti-mouse CD23 PE-Cy7 (B3B4), anti-mouse CD11b A700 (M1/70), anti-mouse CD45R A488 (RA3-6B2), purified anti-mouse CD11b (QA19A45), anti-human CD4 PerCP (RPA-T4), anti-human IgD PE-Cy5 (IA6-2), anti-human CD20 PE-Cy7 (2H7), anti-human IgM A700 (MHM-88), anti-human CD70 FITC (113-16). The following Abs were purchased from Invitrogen: anti-mouse IgD eF450 (11-26c), purified anti-Lipid A (polyclonal), anti-IgM mouse PE-Cy5 (II/41), anti-goat A647 (polyclonal). The following Abs were purchased from eBioscience: anti-mouse CD25 PE-Cy5 (PC61.5) Streptavidin A647 was purchased from BioLegend. CellTrace Violet (CTV) and cytochalasin D was purchased from Invitrogen. Fixable Viability Dye eFluor 506 was purchased from Life Technologies.

### Generation of Phagocytosis Target Beads

Protein conjugated beads were generated using 2 µm fluorescent carboxyl polystyrene beads (FCSY007; Bangs Laboratories) according to the manufacturer’s protocol. In brief, 100 µL of 1% solids bead solution was washed in acetate buffer (0.23% acetic acid, 1% sodium acetate, pH 4.6). After this, beads were resuspended in 10 mg/mL EDAC and were gently mixed for 15 min at room temperature. Beads were washed twice in 1X PBS and resuspended in a solution containing 50 µg/mL of biotinylated OVA (used as target for SAv staining, see below) and one of the following: 85 µg/mL of goat anti-mouse IgM F(ab’)_2_, 85 µg/mL of goat anti-human IgM F(ab’)_2,_ 500 µg/mL of NP-OVA, 500 µg/mL of OVA, 50 µg/mL of mouse C3b, or 25 µg/mL of mouse C3d in 1X PBS and incubated for 2 h at room temperature with constant mixing. Beads were washed and resuspended to a final concentration of 5 × 10^7^ beads/mL in 1X PBS. Beads were stored at 4 °C and used within 4 weeks of production.

Phosphatidylserine-coated beads were generated using 3 μm silica beads (SS05001; Bangs Laboratories) as per our previous work^30^. In brief, 300 µmol of phosphocholine, 80 µmol of phosphoserine, and 40 µmol of rhodamine PE and biotinylated PE in chloroform (78:20:1:1 molar ratio) were added to a small amber vial. The vial was flushed with nitrogen gas to evaporate the lipid mixture. Liposomes were formed by addition of 400 µL of 1X PBS followed by vigorous mixing. 100 µL of 10% solids 3 µm silica bead solution was washed thrice in ddH_2_O, resuspended in the lipid solution, and incubated for 20 min at room temperature with constant mixing. Beads were washed and resuspended to a final concentration of 5 × 10^7^ beads/mL in 1X PBS. Beads were stored at 4 °C and used within 4 weeks of production.

### Mouse Cell Isolation

Lymph nodes and spleens of mice were dissociated in ice-cold EasySep buffer (1 mM EDTA, 2% FBS in PBS). To collect PerC cells, 3 mL of 1X PBS with 0.5% heparin (Sandoz) was injected interperitoneally. An incision in the inner skin of the peritoneum was made to collect the fluid from the cavity. For experiments requiring pure B or T cell populations, B or T cells were isolated using EasySep Negative selection Mouse B, Mouse Pan B, or T cell Enrichment Kits according to the manufacturer’s protocol (STEMCELL Technologies).

### Human B Cell Isolation

Human blood was drawn from healthy donors, and lymphocytes were isolated using Lympholyte-poly Cell Separation Media according to manufacturer’s protocol (Cedarlane). B cells were isolated using EasySep Negative selection Human B cell Enrichment Kits according to the manufacturer’s protocol (STEMCELL Technologies). All experiments performed using human blood was approved by The Western University Health Science Research Ethics Board (2023-123419-82252).

### B Cell Phagocytosis Assays

5 × 10^5^ murine splenic, murine PerC, or human B cells were cultured with 5 × 10^5^ Ag conjugated beads or mRFP1 E. *coli* BL21 in RPMI 1640 containing HEPES (Corning) with 10% FBS and 1% Penstrep for 2 hrs at 37 °C 5% CO_2_. Prior to incubation with beads, some cells were pre-treated with 30 µg/mL cytochalasin D for 1 hr at 37 °C 5% CO_2_. For experiments where Fc receptors or complement receptors were blocked, some cells were pre-treated with 3.85 µg/mL anti-Fcγ receptor, CD16/32 2.4G2 or 1.92 µg/mL anti-mouse CD11b respectively. Cells were subsequently washed, and surface stained for B cell analysis by Flow Cytometry or immunofluorescence microscope.

### Antigen Presentation Assays

Naïve lymph node, splenic, or PerC B cells were isolated from B1-8 Jκ^-/-^ or C57Bl/6 mice, and naïve T cells were isolated from OTII mice respectively. OTII T cells were stained with CellTrace Violet according to the manufacturer’s protocol (Invitrogen). 2.5 × 10^5^cells B cells and 2.5 × 10^5^ OTII T cells were cultured with 5 × 10^5^ Ag conjugated beads, 100 ng/mL OVA, 100 ng/mL NP-OVA, or 5 µg/mL OVA^323-339^ peptide (GenScript) in RPMI 1640 containing HEPES (Corning) with 10% FBS, 1% Penstrep, 1% sodium pyruvate, and 1% non-essential amino acids for 3 or 4 days at 37 °C 5% CO_2_. Cells were subsequently washed, and surface stained for CD4^+^ T cell and B cell analysis by Flow Cytometry.

### Mouse B1 Cell Bacterial Killing Assays

Kanamycin resistant mRFP1 *E. coli* BL21 were received as a gift from Dr. B Heit. Naïve PerC cells were collected from C57Bl/6 mice. 1 × 10^6^whole PerC cells or isolated PerC B cells were cultured with 1 × 10^6^ mRFP1 *E. coli* BL21 in RPMI 1640 with 10% FBS for 4 hrs at 37 °C 5% CO_2_. Cells were subsequently washed and resuspended in RPMI 1640 containing HEPES (Corning) with 10% FBS, 1% Penstrep, 1% sodium pyruvate, and 1% non-essential amino acids and 100 µg/mL of gentamycin (BioShop) and incubated at 37 °C 5% CO_2_ for 1 hr. Cells were then washed and resuspended in RPMI 1640 containing HEPES (Corning) with 10% FBS, 1% Penstrep, 1% sodium pyruvate, and 1% non-essential amino acids and split. Some cells were further incubated for an additional 12 hrs at 37 °C 5% CO_2_. The other cells were washed 3 times in 1X PBS and resuspended in 1X PBS with 0.01% Triton X100 for 10 min at 37 °C 5% CO_2_. Bacteria was then plated on LB agar plates containing 50 µg/mL kanamycin and incubated overnight at 37 °C. After the 12 hr incubation, cells were then washed 3 times in 1X PBS and resuspended in 1X PBS with 0.01% Triton X100 for 10 min at 37 °C 5% CO_2_. Bacteria was then plated on LB agar plates containing 50 µg/mL kanamycin and incubated overnight at 37 °C. CFUs were counted, and bacterial concentrations were calculated to look for evidence of bacterial killing.

### Flow Cytometry

Where indicated, cells were prepared for FACS analysis as previously described^31^. In brief, cells were incubated with anti-Fcγ receptor, CD16/32 2.4G2 (BD Biosciences), in FACS buffer (2% FBS in 1X PBS) for 30 min on ice before further incubation with the indicated Abs. Cells were then stained on ice for 30 min with the Abs listed earlier. Dead cells were identified by staining with the Fixable Viability Dye eFluor506 (eBioscience) according to the manufacturer’s protocol. Cells were washed with FACS buffer between separate stains. Flow Cytometry was performed on a BD Immunocytometry Systems LSRII cytometer and analyzed with FlowJo software (Tree Star).

### Single-Cell RNA-seq (scRNA-seq) Analysis

All scRNA-seq datasets were obtained from the GEO database (https://www.ncbi.nlm.nih.gov/geo/) using the following accession numbers: GSE174739^32^, GSE232834^33^, GSE249975^34^, and GSE210795 (https://www.ncbi.nlm.nih.gov/geo/query/acc.cgi?acc=GSE210795). More specifically, we obtained scRNA-seq of three samples of B cells taken from wild-type (WT) mice, provided by GSE174739. These B cells were taken from the spleen, liver, mesenteric lymph nodes, bone marrow, and peritoneal cavities of mice. Additionally, scRNA-seq data from total peritoneal cells isolated from one sample of WT mice was obtained from GSE232834. Single cell sequencing data from two samples, one from the spleen and one from the peritoneum of WT mice were obtained from GSE249975, and lastly, one dataset of scRNA-seq conducted on total peritoneal cells from macrophage-Cd36 floxed mice was obtained from GSE210795.

Seurat version 5.1.0 was used for the analysis of these scRNA-seq datasets^35^. For each of the scRNA-seq datasets, a Seurat object was created with min.cells and min.features set to 3 and 200, respectively. Cells with unique feature counts greater than 95% or less than 5% quantile were removed. Also, cells that contained a mitochondrial count percentage > 5 or the 90% quantile, whichever was larger, were filtered out. The filtered Seurat objects were merged and normalized using *SCTransform* with default parameters. Seurat *RunPCA* was used to reduce the data dimentionality. This was followed by dataset integration using *CCAIntegration* method. To perform cell clustering, Seurat *FindNeighbors* was run using the first 30 dimensions, followed by running *FindClusters* at a resolution of 0.4.

Marker genes for B1 cells (Ms4a1, Cd19, Ighm, Bhlhe41, Zbtb32), B2 cells (Ms4a1, Cd19, Ighm, Fcer2a, Ighd), and macrophages (Adgre1, Mertk, Fcgr1) were used to identify specific cell clusters which belonged to each of these three cell types. The dataset was then filtered such that only B1 cells, B2 cells, and macrophages remained. The final processed data contained 18,583 genes and 38,676 cells. To identify differentially expressed genes between the three cell types, the *PrepSCTFindMarkers* and *FindMarkers* functions were used. The *FindMarkers* function was run with its recorrect_umi parameter set to false.

### Statistical Analysis

GraphPad PRISM software was used to analyze all FACS data. For statistical comparisons, a one-way ANOVA followed by a t test with Bonferroni correction was used for multiple comparisons. Information about the statistics used for each experiment can be found in the legends. Each data point shown represents a single mouse unless otherwise stated.

## Results

### B cells phagocytose targets via BCR engagement

To visualize BCR-mediated phagocytosis by B cells, we conjugated goat anti-mouse IgM F(ab’)_2_ to 2 µm fluorescent beads. This approach targets unswitched BCR independent of Ag specificity and avoids Fc receptor engagement. Beads were additionally conjugated with biotinylated OVA (OVA_Bio_) to distinguish between extracellular beads and those protected from secondary streptavidin labeling because they have been internalized. Bead targets were incubate at 37 °C 5% CO_2_ with B cells isolated from lymph nodes of GFP^+^ mice and cell:bead interactions were visualized by live fluorescent microscopy. We readily observed B cells engaging with and actively internalizing fluorescent bead targets (**Supplemental Video 1**). In separate experiments, beads were co-cultured with B cells isolated from either lymph nodes or PerC and subsequently stained for cell and bead markers prior to imaging. We observed both streptavidin-labelled external fluorescent bead targets associated with CD19^+^ B cells as well as streptavidin-negative internalized beads within CD19^+^ cells from either tissue source (**Figure 1A**).

**Figure 1:**
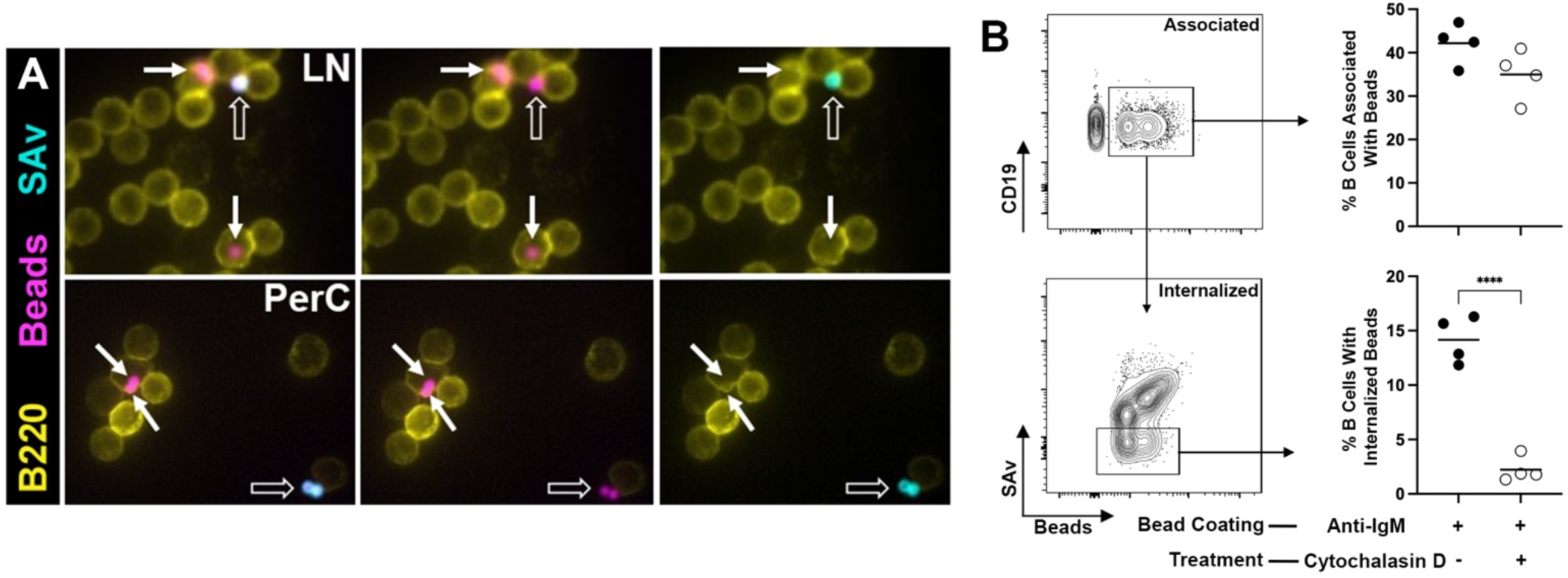
B cells phagocytose targets *via* BCR engagement. Lymph node or peritoneal cavity B cells were isolated from C57Bl/6 and incubated for 2 hrs at 37 °C and 5% CO_2_ with 2 µm fluorescent bead targets with conjugated goat anti-mouse IgM F(ab’)_2_ and OVA_Bio_. (**A**) Representative immunofluorescent microscopy images of LN and peritoneal cavity B cells that are associated with anti-mouse IgM beads, with some remaining extracellular (open arrows) and others internalized (closed arrows). (**B**) Splenic cells were isolated from C57Bl/6 and incubated for 2 hrs at 37 °C and 5% CO_2_ with the bead targets described above. Some cells were pre-treated cytochalasin D for 1 hr at 37 °C and 5% CO_2_. Mean percentage of splenic B cells that were associated with (*top*) or internalized (*bottom*) anti-mouse IgM beads. Each symbol represents data from cells isolated from an individual mouse (n=4). Results from one representative of two independent experiments are shown. ****p < 0.0001 based on a Students t test.

To confirm that we were observing *bona fide* actin-mediated phagocytosis that can be quantified for comparisons between B cell subsets, unisolated C57Bl/6 splenocytes were co-cultured with anti-mouse IgM beads for 2 hrs at 37 °C 5% CO_2_ and then analyzed by Flow Cytometry. Approximately 40% of splenic B220^+^ CD4^-^ B cells co-stained with fluorescent beads (**Figure 1B *top***), and 15% of B cells associated with beads were negative for the secondary streptavidin, indicating that the beads were internalized (**Figure 1B *bottom***). Pre-treatment of cells with cytochalasin D completely prevented internalization of beads but had no impact on association, confirming that we were measuring actin-mediated phagocytosis.

### B1 cells are more phagocytic of BCR targets compared to Fo B and MZ B cells

In the above experiments, phagocytic target beads bound to BCR *via* anti-IgM F(ab’)_2_. To more directly model BCR binding of specific cognate Ag, we developed bead targets coated with Ags recognized by B cells from mice that express knocked-in BCRs with known specificity. To this end, we generated phagocytic targets coated in either NP-haptenated OVA (recognized by B cells from B1-8 Jκ^-/-^ mice) or mMOG_tag_ (recognized by B cells from IgH^MOG^ mice) on beads additionally conjugated with OVA_Bio_ for secondary staining, as described above. Live video microscopy confirmed that isolated lymph node B cells phagocytose cognate Ag beads (**Supplemental Video 2**). Similarly, internalized cognate Ag bead targets were observed in B cells isolated from either the spleen or PerC by immunofluorescence microscopy (**Figure 2A**). Interestingly, B220^dim^ PerC B cells that likely represent B1 cells (see below, **Supplemental Figure 1A**) appeared to internalize more beads that B220^bright^ cells.

**Figure 2:**
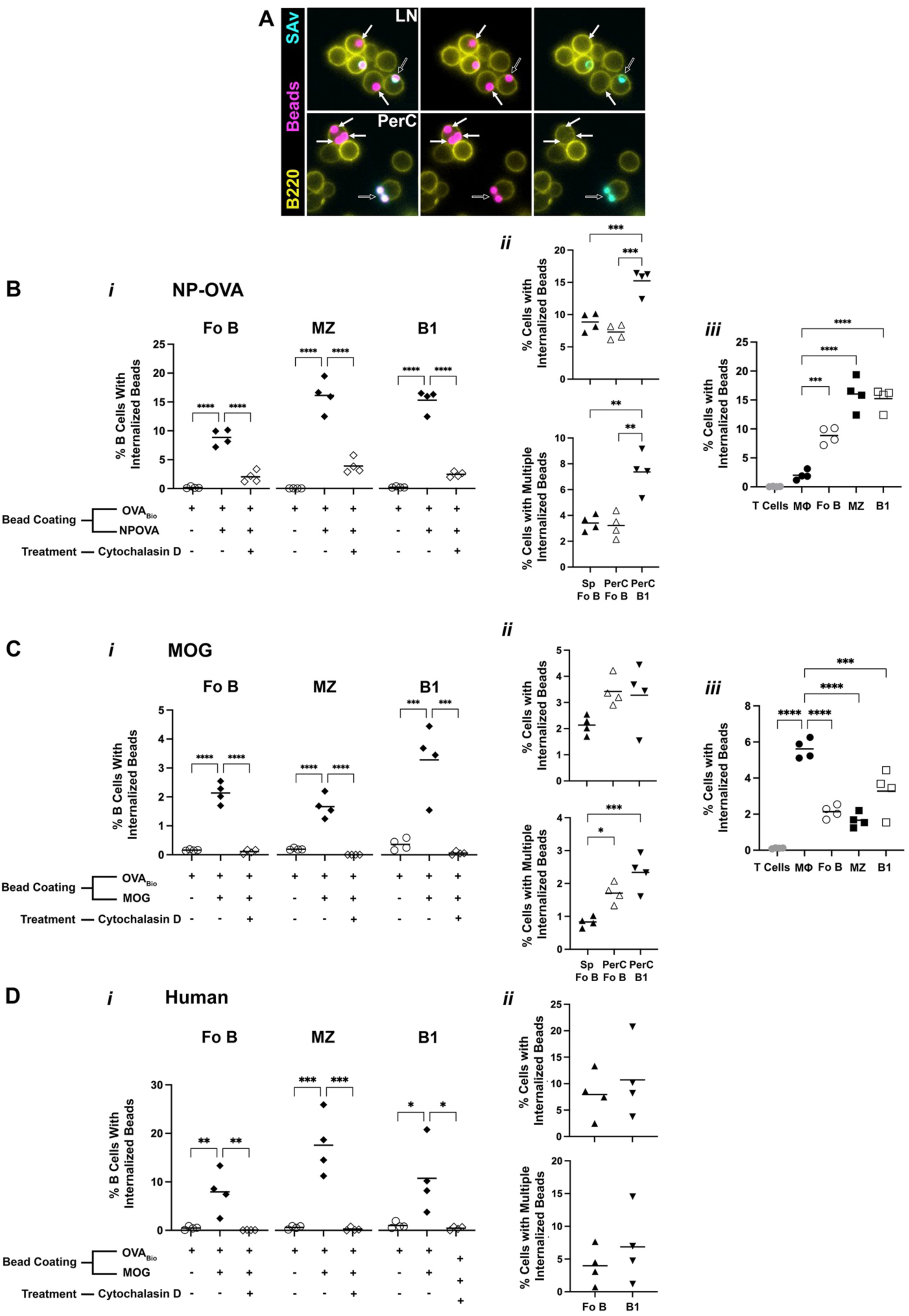
B1 cells are more phagocytic of BCR targets compared to Fo B and MZ B cells. Lymph node or peritoneal cavity cells were isolated from C57Bl/6 and incubated for 2 hrs at 37 °C and 5% CO_2_ with 2 µm fluorescent bead targets conjugated with NP-OVA and OVA_Bio_. (**A**) Representative immunofluorescent microscopy images of splenic and PerC cells with associated (open arrows) or internalized (closed arrows). Splenic and peritoneal cavity cells were isolated from (**B**) B1-8 Jκ^-/-^ or (**C**) IgH^MOG^ and incubated for 2 hrs at 37 °C and 5% CO_2_ with NP-OVA and OVA_Bio_ conjugated beads or MOG and OVA_Bio_ conjugated beads. Some cells were pre-treated cytochalasin D for 1 hr at 37 °C and 5% CO_2_. Mean percentage of Fo B, MZ and B1 cells that internalized (**B*i***) NP-OVA beads or (**C*i***) MOG beads. beads. Mean percentage of splenic Fo B, PerC Fo B, and PerC B1 cells that internalized (**B*ii***) NP-OVA beads or (**C*ii***) MOG beads (*top*), or multiple beads (*bottom*). (*iii*) Mean percentage of T cells, macrophages, and B cells that have internalized (**B*iii***) NP-OVA beads or (**C*iii***) MOG beads. (**D**) Human blood was drawn from healthy female donors and B cells were isolated using EasySep Negative selection Human B Enrichment Kit. Human B cells incubated for 2 hrs at 37 °C and 5% CO_2_ with anti-human IgM beads. Some cells were pre-treated with cytochalasin D for 1 hr at 37 °C and 5% CO_2_. (**D*i***) Mean percentage of Fo B, MZ and B1 like cells with internalized anti-human IgM coated beads. (**D*ii***) Mean percentage of splenic Fo B and B1 like cells that internalized beads (*top*) or multiple beads (*bottom*). In a separate experiment, blood was drawn from healthy male donors, and there were no apparent differences between the two sexes (***data not shown***). Each symbol represents data from cells isolated from an individual mouse or human donor (n=4). Results from one representative of two independent experiments are shown. *p < 0.05, **p < 0.01, ***p < 0.001, ****p < 0.0001 based on an ANOVA followed by a Student t test with Bonferroni correction used for multiple comparisons.

To compare the capacity of Follicular (Fo), Marginal Zone (MZ), and B1 B cell subsets to phagocytose cognate-Ag targets, splenic and PerC cells were collected from B1-8 Jκ^-/-^ or IgH^MOG^ mice which were then incubated with the appropriate cognate Ag bead target or control OVA-coated beads for 2 hrs at 37 °C 5% CO_2_ after which cells were analyzed by Flow Cytometry as described above (see **Supplemental Figure 1B and 1C** for gating).

Splenic Fo and MZ, and PerC B1 subsets did not internalize bead coated with irrelevant OVA protein, but readily internalized beads coated with cognate Ag (either NP-OVA – **Figure 2B*i*** or MOG – **Figure 2C*i***) which was prevented by pre-treatment with cytochalasin D. While the phagocytic capacity of MZ B cells was variable between repeat experiments and between the different bead targets, a consistently greater percentage of B1 cells internalized beads compared to Fo B cells. To confirm that this was not due differences in environment (spleen vs PerC) or cell isolation protocol, we included phenotypic Fo cells from the PerC in a direct comparison between Fo B and B1 cells. Indeed, a greater proportion of PerC B1 cells internalized beads compared to Fo B cells from either the spleen or PerC (**Figure 2B*ii* and C*ii***).

Further, the phagocytic efficiency of PerC B1 cells was greater compared to other subsets, as measured by the ability to internalize more than one bead. This was consistent with our observations by microscopy (**Figure 2A**) where B220^dim^ cells regularly contained multiple internalized beads. Finally, we compared the phagocytic capacity of B cell subsets to other cell types. As expected, T cells did not internalize beads (**Figure 2B*iii* and C*iii***). Interestingly, with respect to NP-OVA-coated targets, all B cell subsets were much superior to professional phagocyte macrophages in internalizing targets that could be bound by BCR (**Figure 2B*iii***). In contrast, IgH^MOG^ B cells were less efficient at internalizing beads coated with the autoantigen MOG, in that a lower percentage of B cells internalized beads (**Figure 2C*iii***). This is expected as, unlike the B1-8 Jκ^-/-^ mice where ∼95% of B cells are specific for NP, only roughly 20% of IgH^MOG^ B cells are specific for MOG. Further, we have previously shown that MOG-binding B cells are less responsive to BCR stimulation than non-specific B cells^36^, which may also impact BCR-mediated phagocytosis.

In similar experiments, we tested the capacity of human B cell subsets to phagocytose BCR targets. Human blood was drawn from healthy donors and isolated human blood B cells were incubated with 2 µm beads coated with goat anti-human IgM F(ab’)_2_ and OVA_Bio_. After 2 hrs of incubation at 37 °C, cells were analyzed by Flow Cytometry as described above (see **Supplemental Figure 1D** for gating). Similar to our observations with mouse cells, circulating B cells with Fo B, MZ, and B1 phenotypes all phagocytosed anti-human IgM beads (**Figure 2D*i***). While there was no consistent difference in phagocytic capacity between human B cell subsets (**Figure 2D*ii***), unlike for mouse B cell subsets, this may be because circulating cells differ from those isolated from the appropriate anatomical niche.

### Fo B cells but not B1 cells efficiently present phagocytosed antigen to T cells

In the context of T cell-dependent antibody responses, Fo B cells internalize Ag bound *via* their BCR to process and present it to activated T cells specific for the same Ag. To measure presentation of phagocytosed Ag to cognate T cells, isolated splenic B cells from B1-8 Jκ^-/-^ mice were co-cultured for 4 d in the presence of target beads and OVA^323-339^ specific CD4 T cells isolated from OTII mice that had been labeled with CellTrace Violet. Proliferation of OVA-specific T cells was used as a measure of the ability of B cells to process whole OVA protein acquired by phagocytosis of the cognate NP-OVA or control OVA bead targets.

B cells cultured with NP-OVA beads induced significantly more proliferation of OTII T cells (**Figure 3A**) and expression of the T cell activation markers CD25 and CD44 (**Figure 3B**) compared to B cells cultured in media alone or with OVA beads. In a separate experiment, we also demonstrated that expression of costimulatory CD86 by B cells is dependent on interactions with T cells, as incubation with Ag alone was not sufficient to upregulate its expression (**Figure 3C**), consistent with the requirement for signals from cognate T cells to activate Ag-engaged Fo B cells that participate in T cell-dependent responses.

**Figure 3:**
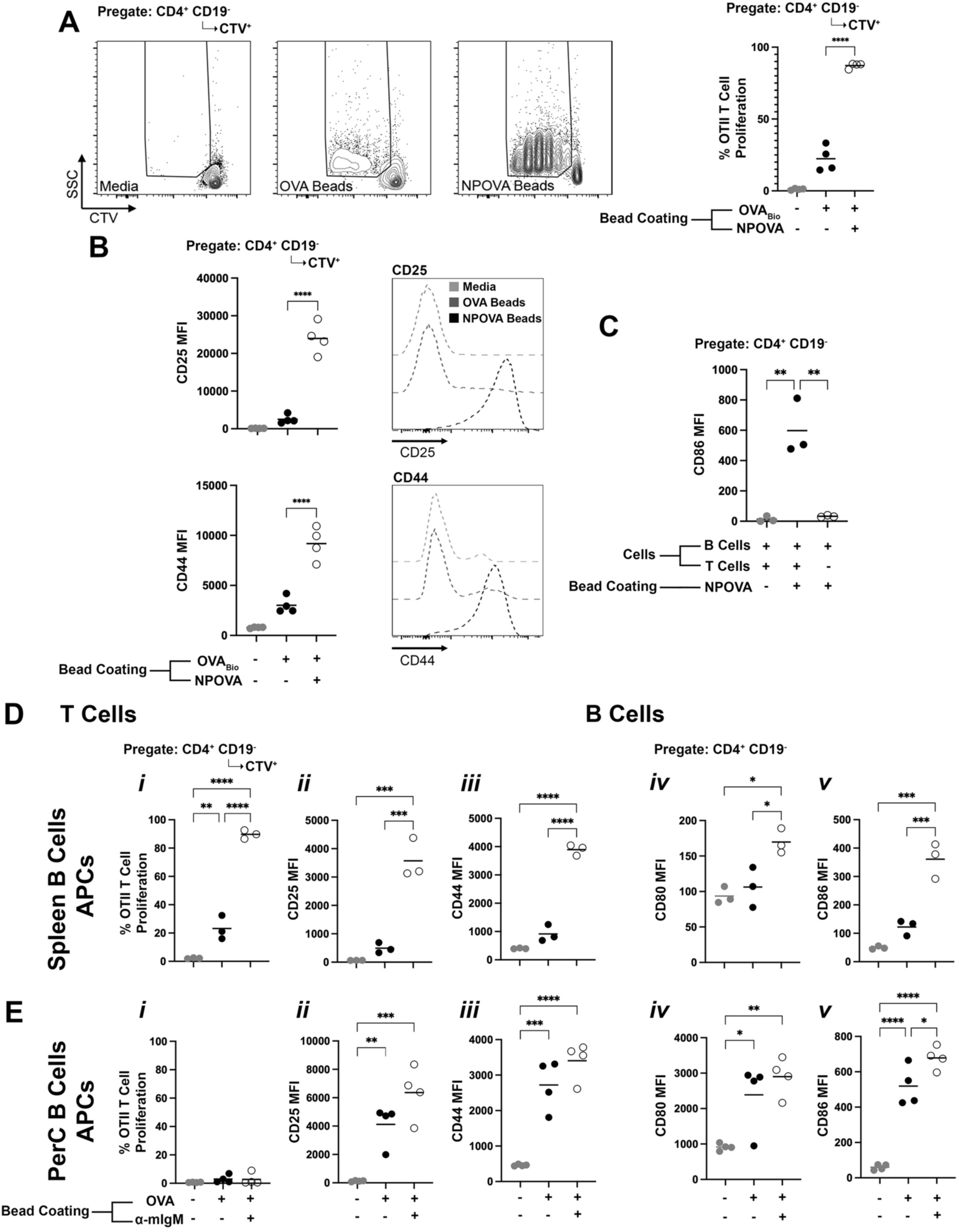
Fo B cells but not B1 cells efficiently present phagocytosed antigen to T cells. Splenic B1-8 Jκ^-/-^ B cells and CellTrace Violet stained CD4^+^ OTII T cells were co-cultured with NP-OVA beads for 4 d at 37 °C and 5% CO_2_. (**A**) Mean percentage of OTII T cells that proliferated. (**B**) Mean MFI of expressed CD25 and CD44 on the surface of CD4^+^ OTII T cells. (**C**) Mean MFI of expressed CD86 on the surface of B1-8 Jκ^-/-^ B cells. (**D**) Splenic or (**E**) peritoneal cavity C57Bl/6 B cells were incubated with CD4^+^ OTII T cells, and anti-mouse IgM and OVA beads for 4 d at 37 °C and 5% CO_2_. (*i*) Mean percentage of CD4^+^ OTII T cells that proliferated and mean MFI of expressed (*ii*) CD25 and (*iii*) CD44 on the surface of CD4^+^ OTII T cells. Mean MFI of expressed (*iv*) CD80 and (*v*) CD86 on the surface of C57Bl/6 splenic or peritoneal cavity B cells. Each symbol represents data from cells isolated from a pair of mice (n=3-4). Results from one representative of two independent experiments are shown. *p < 0.05, **p < 0.01, ***p < 0.001, ****p < 0.0001 based on an ANOVA followed by a Student t test with Bonferroni correction used for multiple comparisons.

B1-8 Jκ^-/-^ mutant mice generate few PerC B1 cells (***data not shown***) and it was not possible to isolate enough of these cells to perform Ag-presentation assays. To compare the Ag-presentation capacity of PerC B1 cells and splenic Fo B cells isolated from wild type C57Bl/6 mice, we made use of the bead targets conjugated to anti-IgM F(ab’)_2_ and OVA_Bio_ protein described above. As observed for the NP-OVA model Ag system, OVA-specific CD4 T cells proliferated significantly more (**Figure 3D*i***) and upregulated the expression of activation markers (**Figure 3D*ii* and *iii***) when co-cultured with splenic B cells in the presence of BCR-targeting anti-IgM/OVA beads compared to media alone or control OVA beads. B cells participating in the cognate interactions similarly upregulated CD80 and CD86 (**Figure 3D*iv* and *v***).

In contrast, B1 cells were unable to support OTII T cell proliferation under any condition (**Figure 3E*i***), even though they did induce significant expression of both CD25 and CD44 on T cells in the presence of both BCR-targeting anti-IgM/OVA beads and, unlike Fo B cells, non-targeting OVA beads (**Figure 3E*ii*, *iii***). The lack of T cell proliferation was not because B1 cells did not express costimulatory molecules CD80 and CD86, which were again upregulated in the presence of BCR-targeting beads and, unlike Fo B cells, non-targeting beads (**Figure 3E*iv*, E*v***).

We performed similar experiments using soluble Ag to determine if the above observation was unique to bead-associated Ag acquired by phagocytosis. B cells were isolated from lymph nodes or the PerC of wild type or B1-8 Jκ^-/-^ mice and cultured for 3 d with CTV-labelled OTII T cells and either no Ag, soluble OVA, soluble NP-OVA, or OVA^323-339^ peptide that corresponds with the OTII TCR epitope and can load directly on cell-surface MHC class II, bypassing the requirement for uptake and processing^37,38^. As expected, wild type C57Bl/6 lymph node B cells did not induce OTII T cell proliferation in the presence of either OVA or NP-OVA protein Ag but did induce proliferation when cultured with OVA^323-339^ peptide (**Figure 4A *top***). Also as expected, lymph node B cells from B1-8 Jκ^-/-^ mice were additionally able to support OTII T cell proliferation when cultured with cognate Ag NP-OVA, but not OVA (**Figure 4A *bottom***). Interestingly, PerC B cells from either C57Bl/6 or B1-8 Jκ^-/-^ B cells only very poorly supported OTII proliferation in the presence of OVA^323-339^ peptide, although both supported a small amount of proliferation in the presence of NP-OVA but not OVA protein. This is consistent with our observations with phagocytic Ag uptake in that B1 cells only poorly supported T cell activation, with some evidence that, unlike Fo B cells, it is not completely limited to Ag acquired by BCR. In separate experiments, we confirmed that Fo B cells and B1 cells expressed similar levels of cell surface MHC class II (**Figure 4B**). Therefore, this does not explain why B1 cells have limited ability to support T cell activation in the presence of protein Ag or pre-processed peptide.

**Figure 4:**
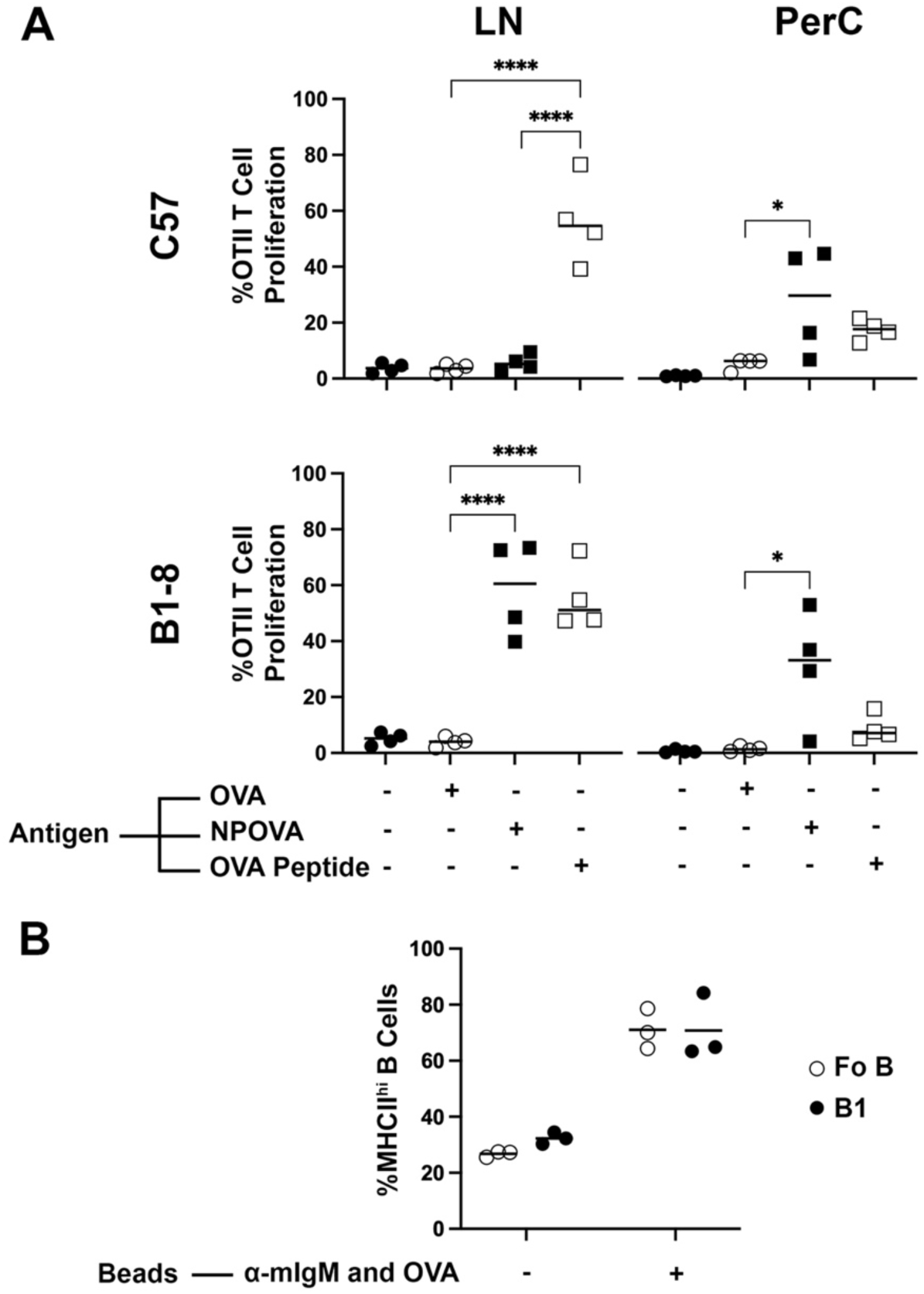
The limited ability of B1 cells to support T cell activation is not due to MHCII expression. Lymph node or PerC B cells from C57Bl/6 or B1-8 Jκ^-/-^ and CellTrace Violet stained CD4^+^ OTII T cells were co-cultured with 100 ng/mL NP-OVA or OVA^323-339^ peptide for 3 d at 37 °C and 5% CO_2_. (**A**) Mean percentage of OTII T cells that proliferated. Splenic C57Bl/6 B cells and CellTrace Violet stained CD4^+^ OTII T cells were co-cultured with anti-mouse IgM and OVA beads for 4 d at 37 °C and 5% CO_2_. Each symbol represents data from cells isolated from a pair of mice (n=4). *p < 0.05, ***p < 0.001, ****p < 0.0001 based on an ANOVA followed by a Student t test with Bonferroni correction used for multiple comparisons. (**B**) Mean percentage of B cells expressing surface MHCII. Each symbol represents data from cells isolated from a pair of mice (n=3). *p < 0.05 based on a two-way ANOVA followed by a Student t test with Bonferroni correction used for multiple comparisons.

### B1 cells, but not other B cell subsets can phagocytose non-cognate antigen

The above results suggest that B1 cells may present Ag to T cells that they are not themselves specific for, albeit infectively. Further, previous studies have suggested that B1 cells can phagocytose bacteria, perhaps recognized via non-BCR mechanisms^24,25^. To identify additional pathways through which B cell subsets may engage phagocytic targets, we modified the beads described above by replacing the BCR-binding elements with opsonins IgG, C3d, or C3b to query Fc Receptor and Complement receptor-mediated phagocytosis. To model efferocytosis of apoptotic cells *via* scavenger receptors, we conjugated silica beads with phosphatidylserine (PS), rhodamine-labelled phosphatidylethanolamine for fluorescence, and biotinylated phosphatidylethanolamine for internalization staining. Splenic and peritoneal cavity cells were collected from C57Bl/6 mice and incubated with the above bead targets for 8 hrs (rather than for 2 hrs, as for the above experiments using BCR-binding beads) followed by analysis by Flow Cytometry (see **Supplemental Figure 2** for gating).

Neither Fo B or MZ B cells internalized IgG, C3b, C3d, or PS beads. In contrast, B1 cells internalized all of the above beads, and this was prevented by pre-treatment with cytochalasin D, confirming actin-mediated uptake (**Figure 5A*i*, B*i*, C*i*, D**). Interestingly, despite our intention to design bead targets to selectively query distinct receptors with known involvement in phagocytosis, blocking antibodies to Fc receptors CD16 and CD32, or complement receptor CD11b had no impact on internalization of IgG or Complement-coated beads, respectively (**Figure 5 A*ii*, B*ii*, C*ii***), likely indicating that B1 cells use multiple, non-selective scavenger receptors to bind to phagocytic targets. Consistent with this, B1 cells internalized the “control” OVA-protein coated beads to the same degree as the targets designed to be recognized by specific receptors (**Figure 5A*i*, B*i*, C*i***). Note that significant internalization of OVA-beads by B1 cells was not evident after 2 hrs of incubation (**Figure 2B*i***), and therefore non-BCR uptake is less efficient compared to BCR-mediated uptake. Indeed, comparison of phagocytic capacity demonstrates that PerC macrophages are substantially better than B cells at internalizing all of the non-BCR targets (**Figure 5 A*iii*, B*iii*, C*iii***), including OVA-protein coated beads (**Figure 5E**).

**Figure 5:**
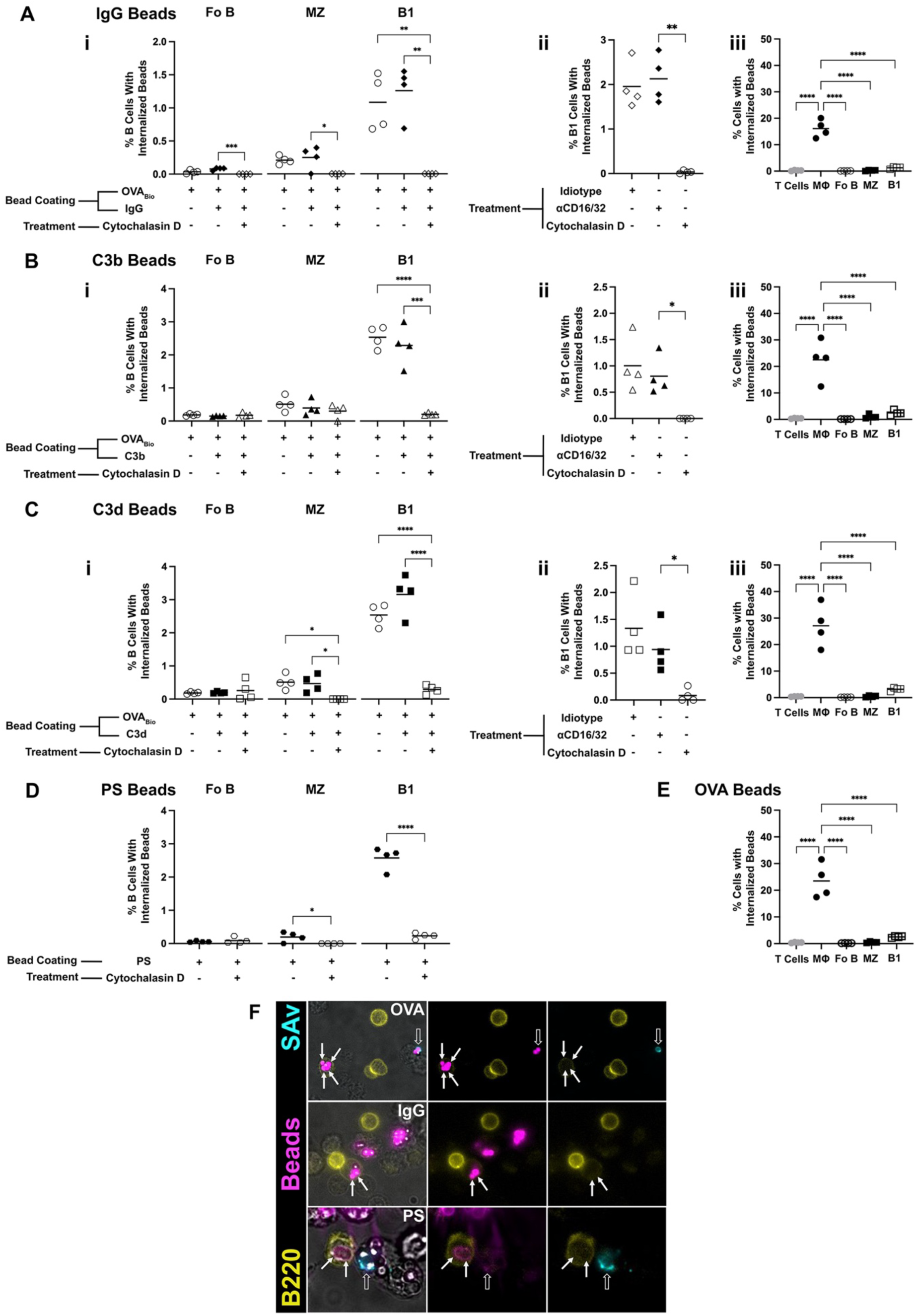
B1 cells, but not other B cell subsets can phagocytose non-cognate antigen. Splenic and peritoneal cavity cells were isolated from C57Bl/6 mice and incubated for 8 hrs at 37 °C and 5% CO_2_ with non-cognate Ag coated beads. Some cells were pre-treated with cytochalasin D, Fc block (anti-CD16/32), anti-CD11b, or idiotype control for 1 hr at 37 °C and 5% CO_2_. Mean percentage of B cells with internalized (**A*i***) IgG, (**B*i***) C3b, (**C*i***) C3d, or (**D**) phosphatidylserine (PS). Mean percentage of PerC B1 cells with internalized (**A*ii***) IgG, (**B*ii***) C3b, or (**C*ii***) C3d beads. Mean percentage of T cells, macrophages, and B cells with internalized (**A*iii***) IgG, (**B*iii***) C3b, (**C*iii***) C3d, or (**E**) OVA beads. (**F**) Immunofluorescence images of peritoneal cavity B cells from a C57/B6 mouse with associated (open arrows) or phagocytosed (closed arrows) OVA, IgG, or PS conjugated beads. Each symbol represents data from cells isolated from an individual mouse (n=4). Results from one representative of two independent experiments are shown. *p < 0.05, **p < 0.01, ***p < 0.001, **** p < 0.0001 based on an ANOVA followed by a Student t test with Bonferroni correction used for multiple comparisons.

PerC B cell internalization of non-BCR-binding beads was confirmed by live video microscopy (**Supplemental Video 3**) and immunofluorescent microscopy (**Figure 5F**). Internalization was largely restricted to B220^low^ B cells that are likely B1 cells, as noted above. Together, these results indicate that, unlike Fo and MZ B cells, B1 cells are not limited to phagocytosing targets bound *via* their BCR. B1 cells are not as efficient at phagocytosing these targets compared to macrophages, nor are B1 cells effective at inducing T cell proliferation in response to presentation of internalized Ag.

### Differences in phagocytic capacity between Fo B cells and B1 cells is not due to differential expression of relevant phagocytic receptors

We have shown that B1 cells can phagocytose multiple targets, while Fo B cells are restricted to BCR-mediated phagocytosis. To determine if this difference in phagocytic capability is due to differential expression of phagocytic receptors, we analyzed public mouse scRNAseq datasets^32,33,34^ (and NCBI accession number GSE210795) containing B cells to identify Fo B cells and B1 cells to compare transcript levels of receptors known to be involved in phagocytosis. As expected, macrophages highly expressed these receptors. B cells also expressed many of the receptors, but there was no apparent difference in expression between Fo B cells and B1 cells (**Figure 6A**). This implies that all B cell subsets may be able to bind targets through non-BCR receptors, but that only B1 cells internalize them. To address this, we analyzed the Flow Cytometry data from the experiments presented in Figure 5 to determine the degree to which different B cell subsets associated with fluorescent bead targets, independent of internalization. Indeed, Fo B cells and B1 cells both bound to IgG, C3b, and C3d beads to similar degrees (**Figure 6B**), but only B1 cells internalized them (**Figure 5A*i*, B*i*. C*i***). Interestingly, MZ B cells bound these beads to an even greater degree, although like Fo B cells they did not internalize them. Therefore, the ability to bind to beads is not the factor limiting subsequent phagocytosis.

**Figure 6:**
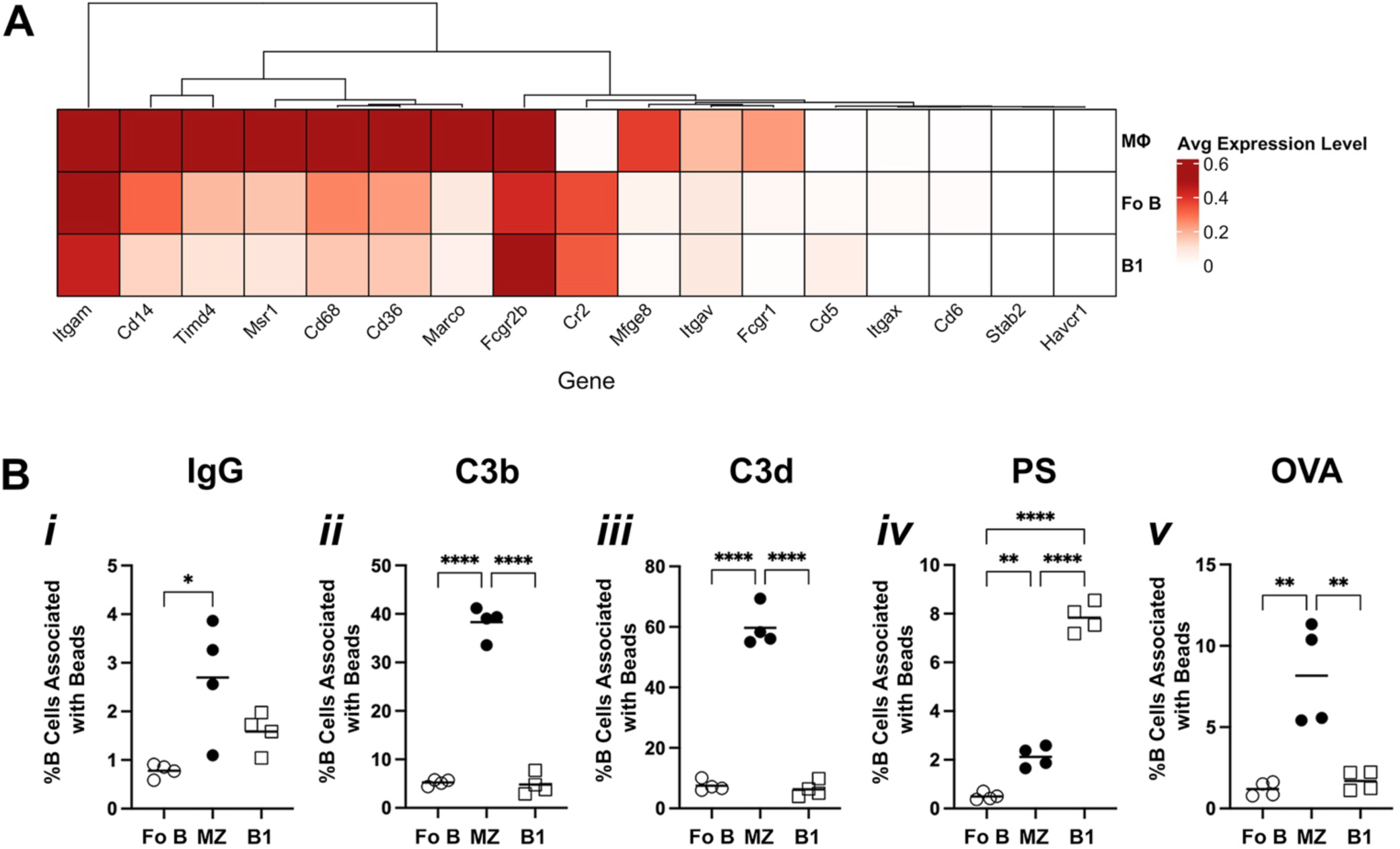
Differences in phagocytic capacity between Fo B cells and B1 cells is not due to differential expression of relevant phagocytic receptors. Publicly available datasets were accessed on NCBI GEO with accession numbers GSE174739, GSE232834, GSE249975 and GSE210795. B2 cells were identified as *Cd19*^+^, *Ms4a1*^+^, and *Fcer2a*^+^. B1 cells were identified as *Cd19*^+^, *Ms4a1*^+^, *Bhlhe41*^+^, and *Fcer2a*^-^. Macrophages were identified as *Adgre1*^+^, *MERTK*^+^, and *Fcgr1A*^+^. (**A**) Average gene expression level of various phagocytic receptors in macrophages, Fo B and B1 cells. Splenic and peritoneal cavity cells were isolated from C57Bl/6 mice and incubated for 8 hrs at 37 °C and 5% CO_2_ with non-cognate Ag coated beads. (**B**) Mean percentage of B cells that were associated with (B*i*) IgG, (B*ii*) C3b, (B*iii*) C3d, (B*iv*) phosphatidylserine, or (B*v*) OVA beads. Each symbol represents data from cells isolated from an individual mouse (n=4). Results from one representative of two independent experiments are shown. *p < 0.05, **p < 0.01, **** p < 0.0001 based on an ANOVA followed by a Student t test with Bonferroni correction used for multiple comparisons.

### B1 cells can phagocytose bacteria but are not effective at killing them

In addition to Ag acquisition, phagocytosis is an important antimicrobial effector mechanism. To determine if B cells kill phagocytosed bacteria, B cells were isolated from the spleen or PerC as described above and incubated with BL21 *E. coli* expressing mRFP1 for 8 hrs at 37 °C 5% CO_2_. Anti-Lipid A labeling was used to differentiate between external and internalized bacteria. By microscopy, we observed that B220^low^ PerC B cells, but not splenic B cells, internalized bacteria (**Figure 7A**). Similarly, analysis of spleen and PerC cells by Flow Cytometry confirmed that PerC B1, splenic Fo B, and splenic MZ B cells bound RFP^+^ bacteria (**Figure 7B**), but only PerC B1 cells could phagocytose them (**Figure 7C**).

**Figure 7:**
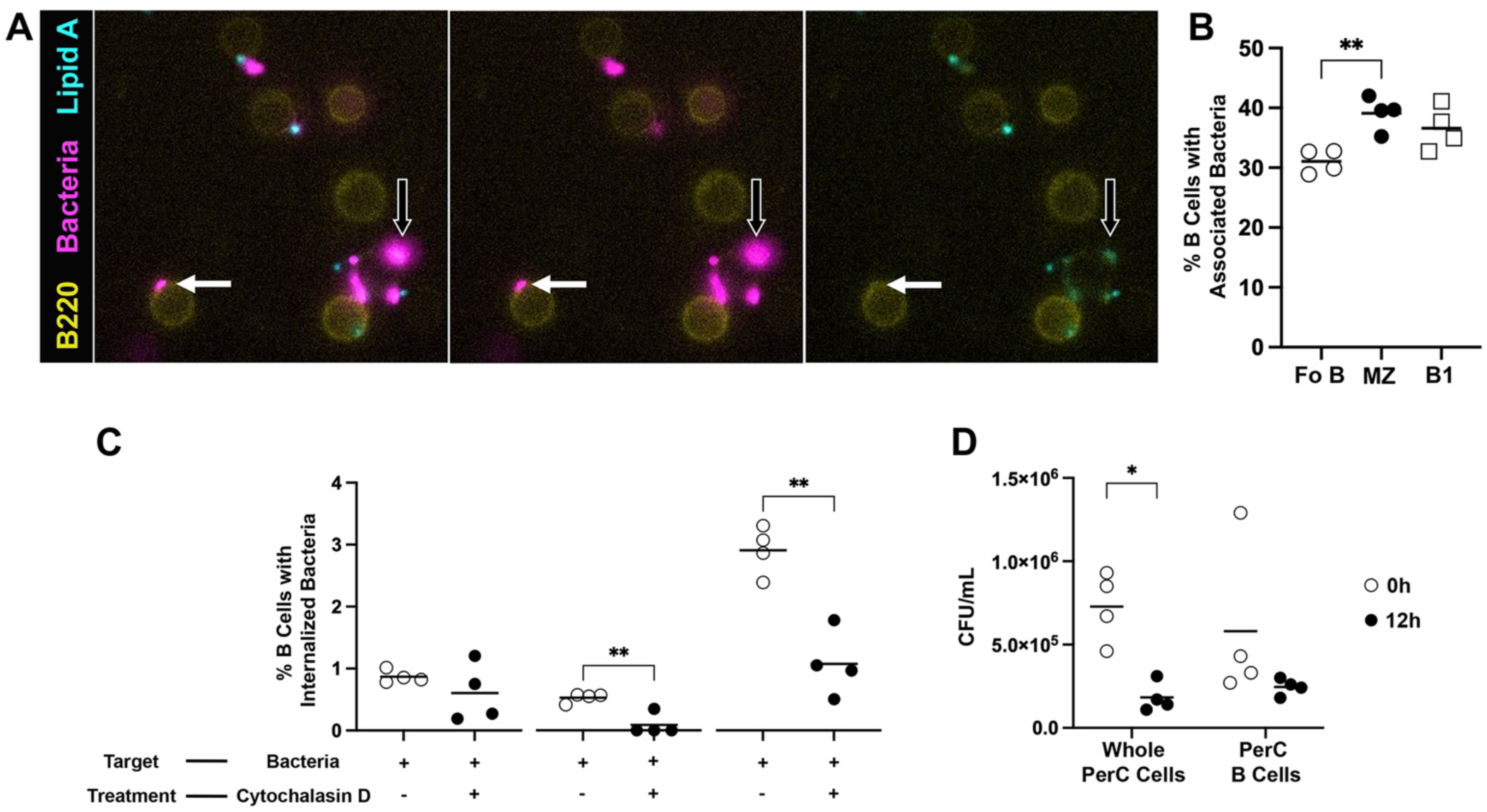
B1 cells can phagocytose bacteria but are not effective at killing them. Splenic and peritoneal cavity cells were collected from C57Bl/6 mice and incubated for 8 hrs at 37 °C and 5% CO_2_ with mRFP1 E. *coli* BL21. Some cells were pre-treated with cytochalasin D for 1 hr at 37 °C and 5% CO_2_. (**A**) Immunofluorescence images of peritoneal cavity B cells from a C57/B6 mouse with associated (open arrows) or phagocytosed mRFP1 E. *coli* BL21 (closed arrows). Peritoneal cavity cells were collected from C57Bl/6 mice and peritoneal cavity B cells were isolated prior to being incubated for 4 hrs at 37 °C and 5% CO_2_ with mRFP1 E. *coli* BL21 followed by gentamycin treatment for 1 hour. Cells were then split, and some cells were permeabilized and internalized bacteria were plated and counted. Some cells were further incubated prior to cell permeabilization and bacterial counting to see if bacteria were killed 12 hrs later. Mean percentage of (**B**) associated with or (**C**) internalized mRFP1 E. *coli* BL21 recovered from PerC cells at the 12 hr timepoint compared with paired PerC cells that were instantly lysed after the 4 hr incubation. Each symbol represents data from cells isolated from an individual mouse (n=4). Results from one representative of two independent experiments are shown. *p < 0.05, **p < 0.01 based on an ANOVA followed by a Student t test with Bonferroni correction used for multiple comparisons.

To determine if B1 cells can kill phagocytosed bacteria, whole PerC cells containing a mixture of macrophages, B1 cells, and other cell types, or isolated PerC B were incubated with mRFP1 *E. coli* BL21 for 4 hrs at 37 °C 5% CO_2_ following by gentamycin treatment for 1 hr to kill any remaining extracellular bacteria. Half of the cells were then lysed to release internalized bacteria for plating on agar (labelled “0 hr”). The other half was incubated for an additional 12 hrs before lysis and plating. The difference in colonies formed between these timepoints represents the killing of phagocytosed bacteria. Indeed, significantly fewer bacteria were recovered from whole PerC cells at the 12 hr timepoint (**Figure 7D**), indicating that some cell type (likely macrophages) effectively killed internalized bacteria. In contrast, there was no difference in the number of bacteria recovered from isolated PerC B cells, suggesting that B1 cells are not efficient bacterial killers.

## Discussion

Here, we performed a survey of the capacity of Fo B, MZ B, and PerC B1 cells to engage with and phagocytose different types of particles by phagocytosis. Our studies provide clarity to previously published reports of B cell phagocytosis using cells from different species and tissue sources^21,22,23,24,25,26,39^. It is not completely clear how B cells gained their reputation of not being phagocytic despite their status as being one of the three professional APCs although we speculate, based on results presented here, that it stems from experiments using Fo B cells and targets that did not engage the BCR^18,19,40^. Regardless it is clear that phagocytosis, as a mechanism to take up potential Ag, is a conserved B cell function across species and subsets. Despite this, phagocytosis uptake is highly regulated in the different B cell subsets, likely commiserate with their different immune roles. Indeed, while all subsets were able to bind different kinds of potential targets, actual internalization through phagocytic mechanisms was restricted to those recognized *via* BCR for Fo and MZ B cells, while only PerC B cells had a broader capacity to internalize *via* other pathways.

A prior study by Parra *et.al.* suggested that PerC B cells but not other B cell subsets are able to phagocytose bacteria engaged through receptors other than the BCR, and our results are in line with these observations^25^. While they did report some bacterial killing by PerC B cells, our results make it clear that B1 cells are not the primary bacterial-killing cells in this anatomical space. How B1 cells recognize bacterial or other non-BCR targets is not entirely clear. Gao *et. al.* postulated that CD11b could be related to the phagocytic ability of B1 cells^24^. However, our results showed that blocking CD11b did not impair uptake of complement coated beads. Our attempts to block the uptake of bead targets designed to engage FcRs also failed. The capacity of PerC B1 cells to phagocytose even apoptotic mimics suggest that they use scavenger receptors with broad specificity, and therefore attempts to block individual receptors were circumvented by redundant recognition systems.

Still, in the absence of robust bacterial killing, the biological significance of B1 cell phagocytosis is not clear. Recent reports have demonstrated that B1 cells can engage in T cell dependent responses^41,42^. Further, Gao *et.al.* reported that PerC B cells could support some T cell activation, especially of already primed T cells^24^. Here, we show that the APC function of B1 cells is very limited, or in fact inhibitory, compared to Fo B cells, an observation that was replicated using both bead-bound and soluble protein Ag, and also short peptide that bypasses internalization and processing. However, B1 cells were similarly capable of expressing MHC class II and costimulatory molecules compared to Fo B cells, and therefore the mechanism(s) limiting interactions between B1 cells and T cells is not clear. The functional consequence may be to uncouple activation and antibody production from T cell dependency, which may be important due to the lack of limits on the pathways used to take up potential Ags by these cells.

In contrast, Fo B cells were highly efficient at presenting Ag acquired by BCR-mediated phagocytosis to cognate T cells, in line with their primary role in T cell-dependent antibody responses where continued interactions with specialized Tfh cells are necessary for B cell activation and subsequent selection of high-affinity clones throughout the GC response^43^. Indeed, Martínez-Riaño *et.al.* demonstrated effective GC formation to particle-bound Ag consistent with phagocytic uptake^26^. Of course, our *ex vivo* presentation assay measures T cell activation by B cells while within the GC it is B cell activation and selection that is dependent on B cell APC function. Nevertheless, as B cell upregulation of costimulatory molecules depend on T cells, it is clear that activation signals travel both directions during the T:B interactions modeled *ex vivo*.

The strict limitation on phagocytic pathways used by Fo B cells is conceptually necessary to protect the integrity of the GC. Uptake and presentation of Ag derived from opsonized targets, for example, would break the link between BCR-specificity and the Ag being presented to Tfh cells. Similarly, uptake and presentation of Ag from apoptotic cells, which are readily available in the GC^44^, could potentially promote autoimmunity. Our results show that block on the use of non-BCR pathways is active, in that murine and human Fo B cells can bind a wide range of targets but only internalize those bound by BCR. However, another study demonstrated that a human B cell line could phagocytose *Mycobacterium tuberculosis*^45^. Non-cognate uptake demonstrated in this study could be explained by the use of a B lymphocyte cell line, which may have a different phenotype compared to primary Fo B lymphocytes.

The mechanism(s) behind the inhibition of uptake *via* non-BCR pathways remains unclear, but functionally Fo B cells bind Ig-opsonized particles *via* the inhibitory FcγRIIb as part of a negative feedback loop to resolve GC responses^46^. Fo B cells can engage with apoptotic bodies through phosphatidylserine receptors such as TIM-1 for the maintenance and induction of IL-10 producing regulatory B cells^47,48^. Further, Fo B cells have been observed to capture proteins and particles non-specifically from the subcapsular sinus of lymph nodes and transport them extracellularly to follicle centres for deposition on Follicular Dendritic Cells^49^. Similarly, MZ B cells which we observed to bind non-BCR targets most readily compared to both Fo B cells and B1 cells, have been reported to transport potential Ags from the red pulp to follicles in the spleen^50,51,52^. Indeed, MZ B cells are identified through their high expression of complement receptors CD21 and CD35^53^. As MZ B cells have been shown to be able engage in T cell dependent responses and form GCs with T cell help^54,55^, the limitation on uptake through non-BCR pathways is likely for similar functional reasons as Fo B cells.

Therefore, Ag uptake machinery in different B cell subsets is tailored to immune function. As “professional” APCs, all B cells express different classes of receptors capable of engaging with different types of particles containing potential Ags and have the capacity to perform phagocytosis in addition to other receptor-mediated mechanism of uptake of smaller particles. However, actual internalization is limited to BCR-bound targets in Fo and MZ B cells, while the ability to bind but not internalize particles may permit the transport of unprocessed Ag extracellularly. For this to be the case, some mechanism must selectively “turn off” non-BCR-mediated phagocytosis in these subsets in mammals, as PerC B1 cells maintain the evolutionarily conserved ability to phagocytose multiple types of targets observed in fish, reptile, and amphibian circulating B cells. How this plays into their biological function is not clear but may be linked to their typical anatomical niche outside of lymphoid spaces.

## Acknowledgements

The authors would like to thank the veterinarians and animal care staff at the West Valley Barrier Facility for their excellent husbandry of our experimental animals. DM was the recipient of an Ontario Graduate Scholarship from the Government of Ontario. This study was funded by an operating grant from the Canadian Institutes of Health Research.

---

**Supplemental Video 1: B cells phagocytose targets *via* BCR engagement.** Lymph node B cells were isolated from a GFP mouse and incubated at at 37 °C and 5% CO_2_ with 2 µm fluorescent bead targets conjugated with goat anti-mouse IgM F(ab’)_2_. Representative live microscopy video of GFP^+^ B cells phagocytosing fluorescent anti-mouse IgM beads.

**Supplemental Video 2: B cells phagocytose cognate antigen coated beads.** Lymph node B cells were isolated from a B1-8 Jκ^-/-^ GFP mouse and incubated at at 37 °C and 5% CO_2_ with 2 µm fluorescent bead targets conjugated with NP-OVA. Representative live microscopy video of NP-specific GFP^+^ B cells phagocytosing fluorescent NP-OVA coated beads.

**Supplemental Video 3: B1 cells can phagocytose non-cognate antigen coated beads.** Peritoneal cavity B cells were isolated from a GFP mouse and incubated at at 37 °C and 5% CO_2_ with 2 µm fluorescent bead targets conjugated with rIgG. Representative live microscopy video of GFP^+^ B cells phagocytosing fluorescent IgG coated beads.

**Supplemental Figure 1:**
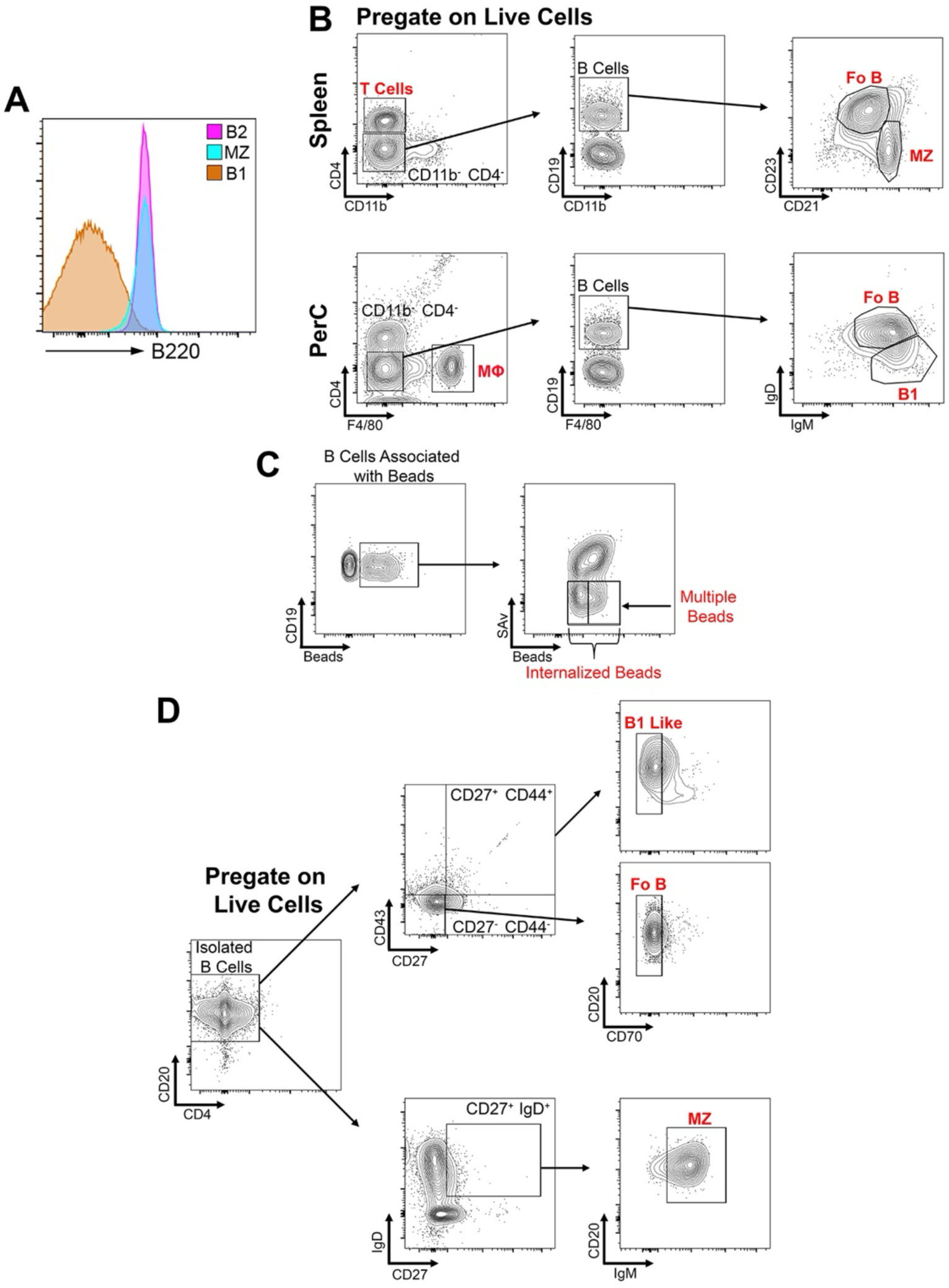
Gating strategy for BCR phagocytosis experiments found in Figure 2. Splenic and peritoneal cavity cells were isolated from B1-8 Jκ^-/-^ or IgH^MOG^ and incubated for 2 hrs at 37 °C and 5% CO_2_ with NP-OVA and OVA_Bio_ conjugated beads or MOG and OVA_Bio_ conjugated beads. (**A**) MFI expression of B220 on the surface of PerC B1, splenic MZ, and splenic Fo B cells. (**B**) Fo B cells and MZ B cells from spleens were identified as CD19^+^, CD4^-^, CD11b^-^, CD21^int^, CD23^+^ and CD19^+^, CD4^-^, CD11b^-^, CD21^hi^, CD23^lo^ respectively. B1 cells were identified from the peritoneal cavity as CD19^+^, CD4^-^, F4/80^-^, IgM^hi^, IgD^lo^. (**C**) After B cell subsets are identified, B cells associated with beads can be identified. Based on SAv staining, we can differentiate the extracellular beads and those protected from secondary streptavidin labeling because they have been internalized. Based on bead fluorescence intensity, we can differentiate B cells that phagocytosed a single bead from B cells that phagocytosed multiple beads. (**D**) Human blood was drawn from healthy female donors and B cells were isolated using EasySep Negative selection Human B Enrichment Kit. Human B cells incubated for 2 hrs at 37 °C and 5% CO_2_ with anti-human IgM beads. Fo B cells were identified as CD20^+^, CD4^-^, CD27^-^, CD43^-^, CD70^-^. MZ B cells were identified as CD20^+^, CD4^-^, CD27^+^, IgD^+^, IgM^int^. B1 like cells were identified as CD20^+^, CD4^-^, CD27^+^, CD43^+^, CD70^-^.

**Supplemental Figure 2:**
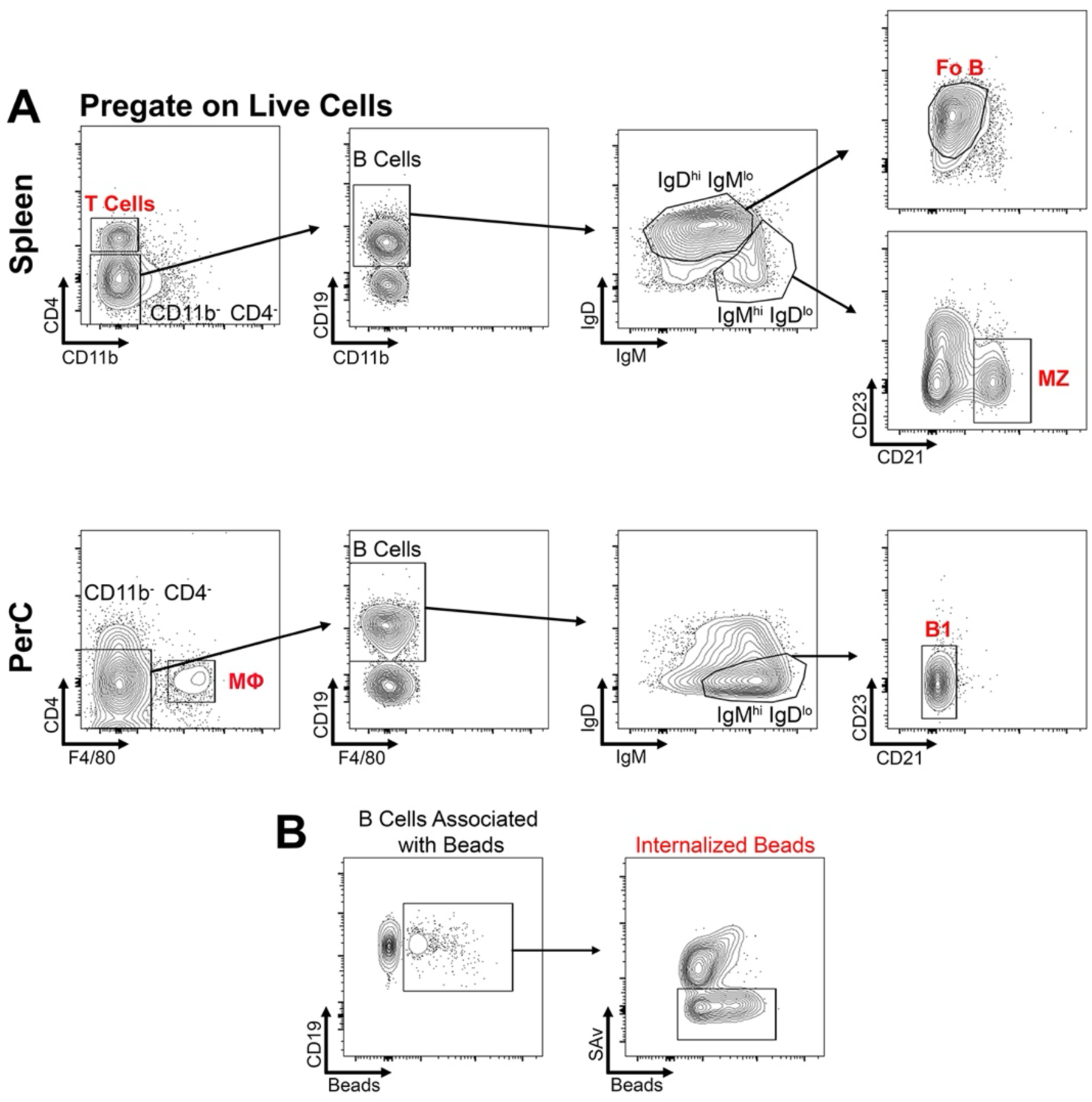
Gating strategy for non-cognate antigen coated bead phagocytosis experiments found in Figure 5. Splenic and peritoneal cavity cells were isolated from C57Bl/6 mice and incubated for 8 hrs at 37 °C and 5% CO_2_ with non-cognate Ag coated beads. (**A**) Splenic Fo B cells and MZ B cells were identified as CD19^+^, CD4^-^, CD11b^-^, IgM^lo^, IgD^hi^, CD21^int^, CD23^+^ and CD19^+^, CD4^-^, CD11b^-^, IgM^hi^, IgD^lo^, CD21^hi^, CD23^lo^ respectively. B1 cells were identified from the peritoneal cavity as CD19^+^, CD4^-^, F4/80^-^, IgM^hi^, IgD^lo^, CD21^-^, CD23^-^. (**B**) After B cell subsets are identified, B cells associated with beads can be identified. Based on SAv staining, we can differentiate the extracellular beads and those protected from secondary streptavidin labeling because they have been internalized. Based on bead fluorescence intensity, we can differentiate B cells that phagocytosed a single bead from B cells that phagocytosed multiple beads.

